# Hot droughts compromise interannual survival across all group sizes in a cooperatively breeding bird

**DOI:** 10.1101/2020.06.11.147538

**Authors:** Amanda R. Bourne, Susan J. Cunningham, Claire N. Spottiswoode, Amanda R. Ridley

## Abstract

Increasingly harsh and unpredictable climate regimes are affecting animal populations around the world as climate change advances. One relatively unexplored aspect of species vulnerability to climate change is whether and to what extent responses to environmental stressors might be mitigated by variation in group size in social species. We used a 15-year dataset for a cooperatively-breeding bird, the southern pied babbler *Turdoides bicolor*, to determine the impact of temperature, rainfall, and group size on body mass change and interannual survival in both juveniles and adults. Hot and dry conditions were associated with reduced juvenile growth, mass loss in adults, and compromised survival between years in both juveniles (−86%) and adults (−60%). Individuals across all group sizes experienced similar effects of climatic conditions. Larger group sizes may not buffer individual group members against the impacts of hot and dry conditions, which are expected to increase in frequency and severity in future.

## Introduction

Anthropogenic climate change is affecting population dynamics across taxa (du Plessis *et al.* 2012; Allen *et al.* 2015; Rey *et al.* 2017; Spooner *et al.* 2018). Understanding life history responses to current environmental conditions is increasingly important for predicting vulnerability to future climate change (Camacho *et al.* 2018; Conradie *et al.* 2019). Species living in arid and semi-arid environments are useful models for studying such responses, because these environments are characterised by extremes in temperature and rainfall (McKechnie *et al.* 2012), and are experiencing rapid increases in temperature and interannual rainfall variability as a result of anthropogenic climate change (Feng & Fu 2013; Mayaud *et al.* 2017). Despite evidence that arid-zone species are well adapted to harsh and unpredictable environments (McKechnie *et al.* 2016; O’Connor *et al.* 2017), increasing temperatures and decreasing rainfall measurably affect behaviour, body condition, growth, and survival in arid-zone birds (McKechnie & Wolf 2010; Cunningham *et al.* 2013; Sunday *et al.* 2014; Iknayan & Beissinger 2018). While droughts are a natural feature of arid and semi-arid ecosystems (MacKellar *et al.* 2014; Tokura *et al.* 2018), an increase in the frequency of ‘hot droughts’ – when above-average temperatures and below-average rainfall co-occur (Overpeck 2013) – is likely (New *et al.* 2006; Kruger & Sekele 2013), with the potential to compromise population persistence in many wildlife species (Walther *et al.* 2002; Sinervo *et al.* 2010; Cruz-McDonnell & Wolf 2016; Paniw *et al.* 2019).

Life-history traits with the potential to mitigate the impacts of high temperatures and drought are of significant interest. Global comparative studies show that the distribution of cooperatively-breeding (Rubenstein & Lovette 2007; Jetz & Rubenstein 2011; Lukas & Clutton-Brock 2017; Shen *et al.* 2017) and group-living (Griesser *et al.* 2017) birds and mammals is associated with harsh and highly variable environments, suggesting that the presence of helpers buffers against environmental uncertainty (Jetz & Rubenstein 2011; Russell 2016; Cornwallis *et al.* 2017), at least up to an optimal number (Markham *et al.* 2015; Ridley 2016). It has been hypothesised that cooperative breeding either evolved in such enviroments (Rubenstein & Lovette 2007; Lukas & Clutton-Brock 2017), enabled species to colonise such environments (Cornwallis *et al.* 2017), or prevented extinction under increasingly harsh conditions (Russell 2016; Griesser *et al.* 2017). One prominent explanation for the occurrence of cooperative breeding in birds is that it represents a ‘bet-hedging’ strategy (Rubenstein 2011): breeding individuals share the costs of reproduction with helpers, enabling them to breed successfully even when conditions are poor (Rubenstein & Lovette 2007). Cooperation may therefore moderate impacts of climate change via task-partitioning (Clutton-Brock *et al.* 2004; Ridley & Raihani 2008), improved access to resources (Golabek *et al.* 2012; Ebensperger *et al.* 2016), or load-lightening (reductions in individual workload in response to the presence of helpers; Crick 1992; Hatchwell 1999; Meade *et al.* 2010; Mumme *et al.* 2015; Langmore *et al.* 2016).

A small number of recent studies empirically test the reproductive benefits of cooperation across varying environmental conditions (Langmore *et al.* 2016; Guindre◻Parker & Rubenstein 2018; van de Ven *et al.* 2020), and one further considers adult survival (Guindre◻Parker & Rubenstein 2020). These empirical tests primarily consider reproduction and survival in response to variation in rainfall. Temperature is rarely included despite the fact that thermoregulatory benefits of group living have been demonstrated (Paquet *et al.* 2016). Behavioural thermoregulation (investing time and energy in self-maintenance, including seeking shade or increasing rest) provides a potential mechanism through which load-lightening may buffer group members against the costs of climate variation. Individuals in larger groups may be able to allocate more time to self-maintenance and therefore suffer fewer consequences of trade-offs between self-maintenance and other essential activities during adverse weather. This may allow higher reproductive success under challenging environmental conditions, and also better mass maintenance and improved survival probabilities.

Long-term monitoring of a population of *Turdoides bicolor* (southern pied babblers, hereafter ‘pied babblers’) provides an opportunity to empirically test the impact of environmental conditions on body mass change (ΔM_b_) and survival between groups of different sizes in a cooperatively breeding species (Ridley 2016). Larger mass is likely to be beneficial in pied babblers because heavier individuals disperse more successfully into breeding positions (Ridley *et al.* 2008). Mass loss occurs when pied babblers provision young (Wiley & Ridley 2016), defend contested territories (Humphries 2013), are evicted from their groups (Ridley *et al.* 2008), or experience high temperatures (du Plessis *et al.* 2012). High temperature extremes have increased in frequency and severity at the study site over the last two decades (van de Ven 2017) and rainfall is extremely variable from year to year (MacKellar *et al.* 2014). In pied babblers, high temperatures and/or drought increase the risk of local extinction (Wiley 2017), reduce offspring provisioning rates (resulting in smaller nestlings; Wiley & Ridley 2016), limit foraging efficiency (du Plessis *et al.* 2012), lower daily energy expenditure (Bourne *et al.* 2019), and decrease investment in territorial defence (Golabek *et al.* 2012).

Here, we examine how within-season ΔM_b_ and interannual survival in pied babblers varies with temperature, rainfall, and group size. We expected negative effects of high temperatures, and positive effects of high rainfall and larger group sizes, on 1) ΔM_b_ in juveniles and in breeding adults, 2) survival of juvenile birds from nutritional independence at 90 days of age to recruitment into the adult population at one year of age, and 3) survival of breeding adults from one breeding season to the next. Analyses do not include subordinate adults because subordinate adults (both sexes) often disperse (Ridley *et al.* 2008; Raihani *et al.* 2010) and dispersal is easily confounded with mortality (Layton-Matthews, Ozgul, & Griesser, 2018). We further predicted that larger group sizes would buffer against climatic effects due to load-lightening allowing individuals to invest more in self-directed behaviours during periods of harsh weather. Specifically, we predicted that individuals in larger groups would experience disproportionately fewer negative effects of high temperatures and drought on both ΔM_b_ and survival.

## Materials and methods

### Study site and system

Pied babblers are medium-sized (60–90 g), cooperatively-breeding passerines endemic to the Kalahari Desert in southern Africa (Hockey *et al.* 2005). Fieldwork was undertaken at the 33 km^2^ Kuruman River Reserve (KRR; 26°58’S, 21°49’E). Mean summer daily maximum temperature in the region averages 34.7 ± 9.7°C, and mean annual precipitation averages 186.2 ± 87.5mm (1995–2015, van de Ven, McKechnie & Cunningham 2019). Rainfall has been declining and high temperature extremes increasing in both frequency and severity over the last 20 years (Kruger & Sekele 2013; van Wilgen *et al.* 2016; van de Ven 2017).

Pied babbler group sizes range between 3–15 adults (Ridley 2016). Groups consist of a single breeding pair, one or more subordinate adult helpers, and immature offspring (Nelson-Flower *et al.* 2011). All adult group members cooperate, participating in territorial defence, sentinel behaviour, and caring for young (Ridley 2016). Pied babblers are considered nutritionally independent (receiving < 1 feed per hour; referred to as ‘juveniles’) by 90 days of age (Ridley & Raihani 2007), and are defined as sexually mature adults one year after hatching (Ridley 2016).

Birds in the study population are habituated to observation at distances of 1–5 m (Ridley & Raihani 2007). Group composition and life history checks are conducted weekly with habituated groups throughout each summer breeding season (September to March). Groups are territorial and can be reliably located by visits to each territory. Birds in the study population are marked with a unique combination of metal and colour rings for individual identification.

### Data collection

Data were collected for each breeding season between September 2005 and 2019. Pied babblers are sexually monomorphic (Ridley 2016; Bourne *et al.* 2018) and molecular sexing was used to determine sex of individuals (following Fridolfsson & Ellegren, 1999). Blood samples for sexing were collected by brachial venipuncture.

#### Body mass measurements

Nestlings were weighed to 0.1 g on a top-pan scale 11 days post-hatching (Mass_11_). Body mass data were collected from 90 (± 15) day-old birds (Mass_90_; *n* = 323 mass measurements from 124 individuals) by enticing individuals to stand on a top-pan balance in exchange for a small food reward (Ridley 2016). Most measurements (74%) were collected within 7 days of the 90 day mark. Mean mass was calcultated where multiple measurements per individual were available (52% of juveniles). The sampling period is justified because pied babblers are approximately fully-grown by 90 days of age (Raihani & Ridley 2007; Thompson et al. 2013; Ridley 2016). This was confirmed by data from 16 individuals from whom we collected ≥ 5 mass measurements between 75 and 105 days of age [1 (6%) maintained mass over the period, 7 (44%) lost mass, and 8 (50%) gained mass, with an average change in mass between earliest and latest measurement of ~1 g].

Body mass data were collected from adult breeding birds from the beginning (September or October, Mass_Oct_) and end (February or March, Mass_Mar_) of each breeding season in the same way as for juveniles. Variation in exact sampling periods for each mass measure reflect annual variation in the timing of the breeding season. We chose the difference between Mass_Oct_ and Mass_Mar_ as a biologically relevant measure because this time period encompasses a) the hottest months at the study site (December, average T_max_ = 35.71°C; and January, average T_max_ = 35.69°C), b) the wettest months at the study site (72% of annual rain falls December-February), and c) the core breeding season (68% of breeding occurs October-December). We collected multiple Mass_Oct_ and Mass_Mar_ measurements for 63% and 52% of the birds respectively, and used the average of all mass measurements per individual where available.

All body mass measurements from juveniles and adults were collected at dawn, representing pre-foraging body mass. Body mass change (ΔM_b_) was calculated in grams as follows: ΔM_b_ between 11 days and 90 days of age for juveniles (ΔM_b.Juv_) = Mass_90_ − Mass_11_; and ΔM_b_ from the start to the end of the breeding season for breeding adults (ΔM_b.Adults_) = Mass_Mar_ − Mass_Oct_.

#### Life history data

##### (i) Juvenile birds

All nests initiated in the study population during each breeding season were monitored to determine hatching dates. Each brood was checked daily from 14 days post-hatching onwards, to determine the date on which nestlings fledged. Natal group size (G.Size_Brood_; the number of adults present in each individual’s natal group between hatching and fledging; constant during the chick rearing period for all but five of 147 nests), average group size for the period between fledging and independence (G.Size_90;_ variations in group size due to older juveniles reaching 1 year of age, or dispersal of subordinate adults, were observed in 58 of 147 breeding attempts), and brood size (number of nestlings in the brood 11 days post-hatch) were recorded for each brood. Presence/absence of fledglings was noted during weekly visits within each breeding season and presence/absence of juvenile pied babblers was recorded at one year (±15 days) post hatching. Presence/absence in the population at one year of age represents a ‘disappearance rate’ likely to be driven primarily by mortality. Dispersal typically occurs after individuals have reached sexual maturity and the mean age at first dispersal is 565 days (Nelson-Flower *et al.* 2018). Individuals that dispersed before one year of age (*n* = 1) were excluded from the analysis.

##### (ii) Adult breeding birds

Dominant individuals can be identified unambiguously through incubation behaviour (Ridley 2016) and distinctive duets (Wiley & Ridley 2018). Presence/absence of dominant individuals was recorded at the beginning of each breeding season during an annual census, at which point we could determine whether breeding adults had survived over the most recent winter putting them in the position to breed again. For breeding adults, territory and pair fidelity is high between years and voluntary dispersal is unusual (Raihani *et al.* 2010; Wiley & Ridley 2018). Overwinter disappearance is therefore likely to be driven primarily by mortality. Data for individuals who were dominant for only part of a breeding season due to death (*n* = 47) or dispersal (*n* = 9) were excluded from analyses of interannual survival. We calculated average group size during each breeding season (G.Size_BrSeas_; variations in group size were observed in 62 of 177 group-seasons).

#### Temperature and rainfall

Daily maximum temperature (°C) and rainfall (mm) data were collected from an on-site weather station (Vantage Pro2, Davis Instruments, Hayward, USA; see van de Ven, McKechnie, & Cunningham, 2019). Missing data from 2009, 2010, and 2011 were sourced from a nearby South Africa Weather Services (SAWS) weather station (Van Zylsrus, 28 km), which recorded significantly repeatable temperature measurements (Lin’s concordance correlation coefficient *r_c_* = 0.957, 95 % CI: 0.951–0.962), and moderately repeatable rainfall measurements (*r_c_* = 0.517, 95% CI: 0.465–0.566) in comparison with the on-site weather station. Differences in rainfall were small (average difference = 0.045 ± 3.075 mm, 95 % CI = −5.981–6.072 mm), suggesting that both weather stations adequately detected wet vs. dry periods.

For analyses relating to juveniles, daily maximum temperatures were averaged for the offspring developmental periods between hatching and fledging (mean T_maxBrood_) and between fledging and independence (mean T_max90_). Rainfall was summed for the 60 days prior to initiation of incubation at the nest from which each individual fledged (Rain_60_; to allow for delays between rainfall and invertebrate emergence; Cumming & Bernard, 1997; Ridley & Child, 2009) and for the period between fledging and independence (Rain_90_). For analyses relating to adults, daily maximum temperatures were averaged, and rainfall was summed, over the whole breeding season (Sept – Mar, Mean T_maxBrSeas_; Rain_BrSeas_). Long term rainfall data for the region was used to determine the presence or absence of a meteorological drought within a breeding season (Drought_BrSeas_). These data were obtained from a SAWS weather station at Twee Rivieren (~120 km from the study site; available until 2013). Following Mayaud et al (2017), meteorological drought was defined as ≤ 75% of average precipitation between September and March (≤ 137.75 mm), using long term data for the 30-year period 1984–2013 to determine average precipitation in the region.

###### Box 1: Glossary of terms

Brood size: Number of nestlings per brood
G.Size_Brood_: Number of adults in the natal group at initiation of incubation
G.Size_90_: Average number of adults in the natal group between fledge and nutritional independence at 90 days of age
G.Size_BrSeas_: Average number of adults in the natal group over the breeding season
Mass_11_: Nestling body mass (0.1 g) collected 11 days after hatching
Mass_90_: (Average) body mass data (0.1 g) collected from 90 (± 15) day-old birds
ΔM_b.Juv_: Change in juvenile body mass (g), calculated as Mass_90_ − Mass_11_
Mass_Oct_: (Average) body mass data (0.1 g) collected from adults at the beginning of the breeding season (September and October)
Mass_Mar_: (Average) body data (0.1 g) collected from adults at the end of the breeding season (February and March)
ΔM_b.Adults_: Change in adult body mass (g), calculated as Mass_Mar_ – Mass_Oct_
Mean T_maxBrood_: Average daily maximum temperatures between hatching and fledging
Mean T_max90_: Average daily maximum temperatures between fledging and nutritional independence at 90 days of age
Mean T_maxBrSeas_: Average daily maximum temperatures between the start (September) and the end (March) of the breeding season
Rain_60_: Total rainfall in the 60 days prior to initiation of incubation at the nest from which each individual fledged
Rain_90_: Total rainfall between fledging and nutritional independence at 90 days of age
Rain_BrSeas_: Total rainfall between the start (September) and the end (March) of the breeding season
Drought_BrSeas_: Occurrence of a meteorological drought (rainfall < 135.75 mm) within the breeding season

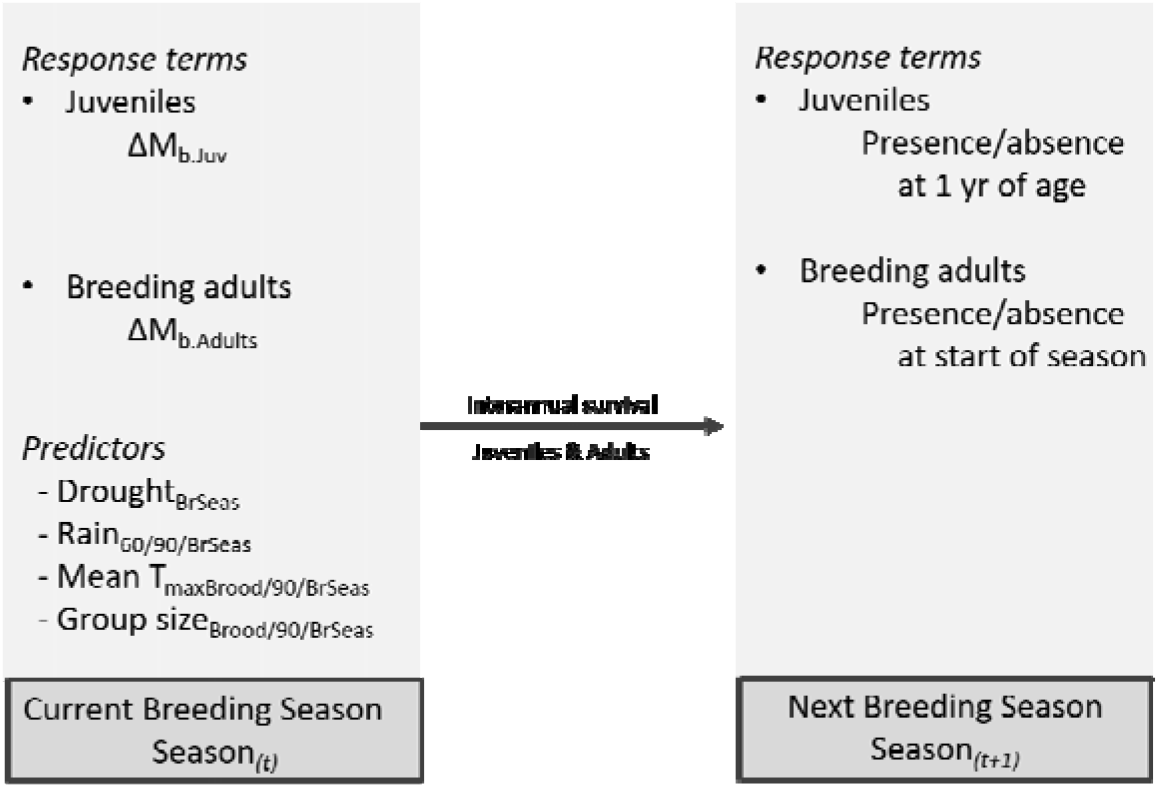
Caption: The grey boxes below show the breeding seasons within which data were collected. All the climate-related predictor variables were collected during Breeding Season_(t)_ and used in models testing variation in body mass change within the same Breeding Season_(t)_ and interannual survival into the following Breeding Season_(t+1)_

### Statistical analyses

Statistical analyses were conducted in R v 3.4.1 (R Core Team 2017). Mixed effects models, using the package *lme4* (Bates *et al.* 2015), were used for all analyses, see Box 1. Model selection with Akaike’s information criterion corrected for small sample size (AICc) was used to explore a series of models to determine which parameters best explained patterns of variation in the data (Symonds & Moussalli 2011; Harrison *et al.* 2018). Where multiple models were within 5 AICc of the top model, top model sets were averaged using the package *MuMin* (Barton 2015) and we present parameter estimates for interpretation after model averaging (Burnham & Anderson 2002; Grueber *et al.* 2011). All continuous explanatory variables were scaled by centering and standardising by the mean (Schielzeth 2010; Harrison *et al.* 2018). All explanatory variables were tested for correlation with one another (all VIF < 2 for pairs of continuous predictors, Fox & Monette 1992). Rainfall measures were always highly correlated with drought (*F*_1,247-350_ > 37.611, *p* < 0.001), and therefore rainfall and drought were not included in the same additive models (Harrison *et al.* 2018). Linear model fits were checked using normal Q-Q plots and histograms of residuals. Binomial model fits were checked for overdispersion in the *RVAideMemoire* package (Herve 2019) and for zero-inflation in the *DHARMa* package (Hartig 2020). Model terms with confidence intervals not intersecting zero were considered to explain significant patterns in the data (Grueber, Nakagawa, Laws, & Jamieson, 2011). Sample sizes reflect datasets after removing records containing missing values. Unless otherwise indicated, summary statistics are presented as mean ± one standard deviation.

Interactions between group size and climatic effects on ΔM_b_ and survival would be consistent with a buffering effect of group size. We conducted sensitivity power analyses to identify the minimum determinable effect of two-way interactions given our sample sizes (Cohen 1988; Greenland *et al.* 2016). Assuming a fourfold increase in required sample size to adequately detect interactions in mixed-effects linear regression models (Leon & Heo 2009), we confirmed sufficient sample size to detect 1) small to moderate main effects in all analyses (Cohen’s *f^2^* ≤ 0.12 in all cases), 2) moderate to very large effects of two-way interactions in ΔM_b_ analyses (*f^2^* = 0.28 for fledglings, *f^2^* = 0.54 for adults), and 3) small to moderate effects of two-way interactions in interannual survival analyses (*f^2^* ≤ 0.14 for juveniles, *f^2^* = 0.09 for breeding adults).

Where interannual survival probabilities for juveniles and breeding adults were influenced by interactions, we used the package *lsmeans* (Lenth 2016) to predict survival probabilities based on different values of the interacting factors.

#### Body mass change

To determine which variables explained ΔM_b.Juv_ and ΔM_b.Adults_, we used maximum likelihood linear mixed-effects models (LMMs). For ΔM_b.Juv_ (*n* = 124), we considered the influence of G.Size_90_, Drought_BrSeas_, Rain_60_, Rain_90_, mean T_max90_, sex, brood size, and the interactions among climate variables and group size. Brood identity nested within group identity were included as random terms to account for repeated measures. For ΔM_b.Adults_ (*n* = 82 measurements from 53 different individuals), we considered the influence G.Size_BrSeas_, Drought_BrSeas_, Rain_BrSeas_, mean T_maxBrSeas_, age (in days since hatching), sex, and the interactions among climate variables and group size. Individual identity nested within group identity were included as random terms.

#### Survival

Non-monitoring periods over winter prevented detailed time-step survival analyses, such as Cox proportional hazards models (Cox 1972; Austin 2017; Guindre◻Parker & Rubenstein 2020), for both juveniles and adults. Therefore, to determine which variables explained interannual survival of known individuals in the study population, we used generalised linear mixed-effects models (GLMMs) with a binomial distribution (survival to next breeding season = yes/no) and a logit link function.

*Juveniles:* For juvenile birds, interannual survival was measured as survival from nutritional independence to one year of age (± 15 days; recorded in the following breeding season – see Box 1). The factors that influence pied babbler survival probabilities are not constant across time during early development (Bourne *et al.* n.d.; Ridley 2016), and conditions experienced in the nest can carry over to influence survival probabilities later (Harrison *et al.* 2011; Auer & Martin 2017; Moore & Martin 2019). We therefore conducted separate analyses specifically considering climate and social factors experienced in the nest and after fledging. For survival to one year, we considered the influence of (a) G.Size_Brood_, Rain_60_, Drought_BrSeas_, mean T_maxBrood_, sex, Mass_11_, brood size, and the two way interactions amongst all group size and climate variables (i.e conditions between hatching and fledging; *n* = 247 individuals); and (b) G.Size_90_, Rain_90_, Drought_BrSeas_, mean T_max90_, sex, Mass_11_, brood size, and the two way interactions among all group size and climate variables (i.e. conditions between fledging and nutritional independence; *n* = 229 individuals). Brood identity nested within natal group identity were included as random terms in both analyses.
*Breeding adults:* For breeding adults, interannual survival was measured from the end of a breeding season in which they attempted to breed to the beginning of the subsequent breeding season (see Box 1). For interannual survival (*n* = 352 records from 136 different adults), we considered the influence of G.Size_BrSeas_, Drought_BrSeas_, Rain_BrSeas_, Mean T_maxBrSeas_, age, sex, and the two way interactions among all group size and climate variables. Individual identity nested within group identity were included as random terms.

We tested for the influence of ΔM_b_ on interannual survival probabilities for both age classes separately, using univariate binomial GLMMs with a logit link function, due to much smaller sample sizes for body mass than for presence/absence data.

## Results

Annual average summer maximum temperature at the study site from 2005–2019 was 34.2 ± 0.9 °C (range: 32.4–36.5 °C), summer rainfall averaged 185.4 ± 86.2 mm (range: 64.4– 352.1 mm), and droughts occurred in 5 of 14 breeding seasons studied (Fig. 1A). Group size varied between groups and breeding seasons, averaging 4.2 ± 1.4 adults per group across all breeding seasons (range: 2–9 adults; Fig. 1B), but did not differ significantly between drought and not-drought years (*F_1,164_* = 0.754, *p* = 0.387). Between 2005 and 2019, the largest group averaged 5.4 ± 2.3 adult group members (range across 11 breeding seasons: 2.3–9), while the smallest group averaged 3.3 ± 0.9 members (range across 12 breeding seasons: 2–5).

**Fig 1:**
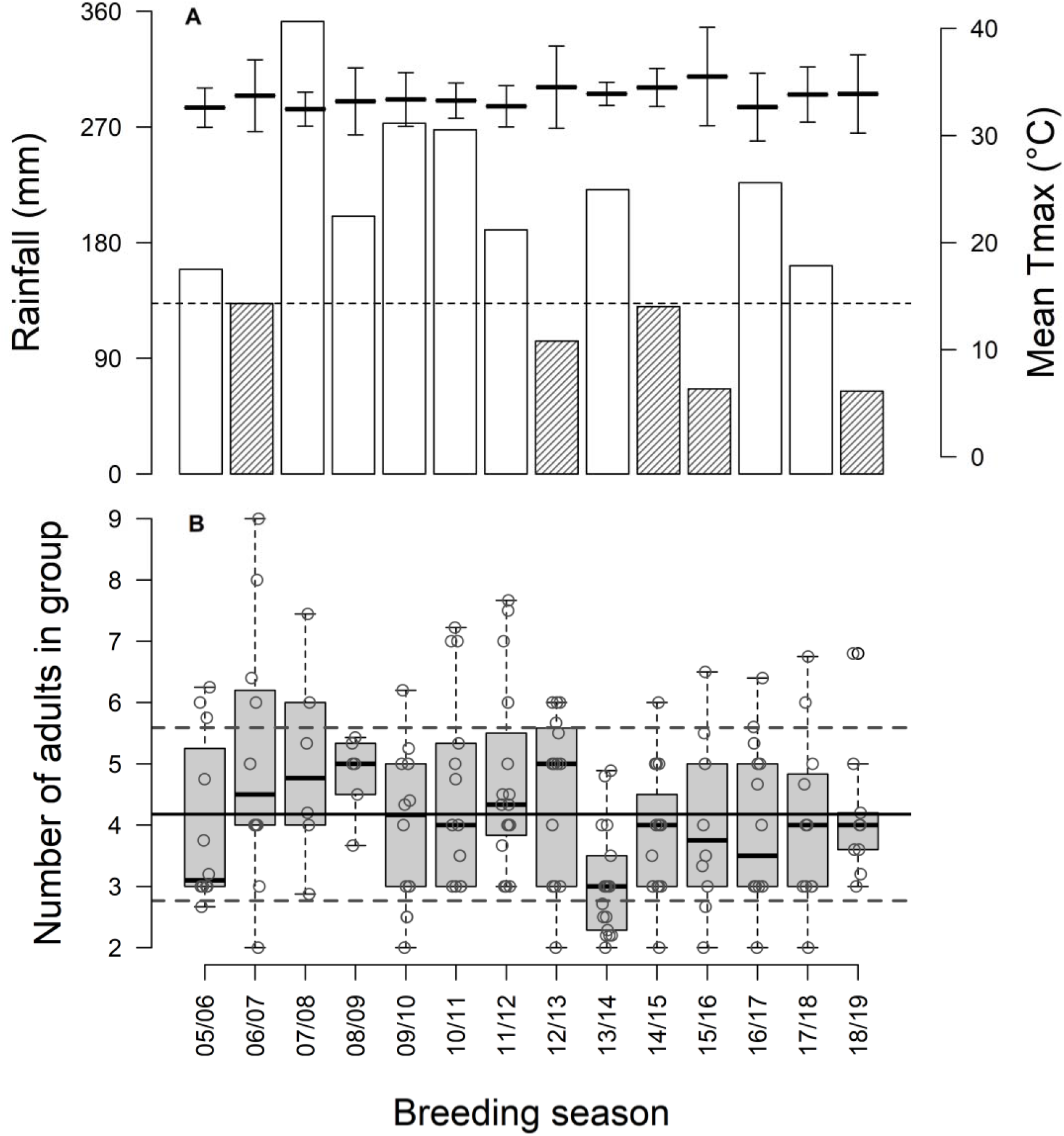
(A) Average maximum temperature (black dashes ± 1 SD) and total rainfall (vertical bars: no colour = no drought, hatched = drought) for each austral summer breeding season studied from 2005 to 2019. Dashed horizontal line represents rainfall = 137.75 mm; years with rainfall < 137.75 mm were classified as drought years. (B) Boxplots show median (black line), first and third quartiles (box), and interquartile range (whiskers) for the distribution of average group size across each breeding season. Open circles represent data points (jittered for improved visibility) and lines present the study-wide average group size (solid horizontal line) ± 1 SD (dashed horizontal lines).

### Change in body mass

Fledglings that survived to 90 days were heavier as nestlings (mean Mass_11_ = 40.0 ± 5.5 g, *n* = 270) than those that did not survive (37.4 ± 7.3 g, *n* = 295; LMM with brood identity as the random term: Est = 2.403 ± 0.526, *t* = 4.566, *95% CI* 1.371, 3.437). Individuals gained significantly more mass between fledging and independence during wetter periods, and when they were raised in larger broods (Table 1A, Fig. 2A, Fig. 2B). ΔM_b.Juv_ did not vary significantly with sex, group size or temperature between fledging and independence, nor did we find evidence of significant interactions among climate and group size variables (Table S1).

**Table 1:**
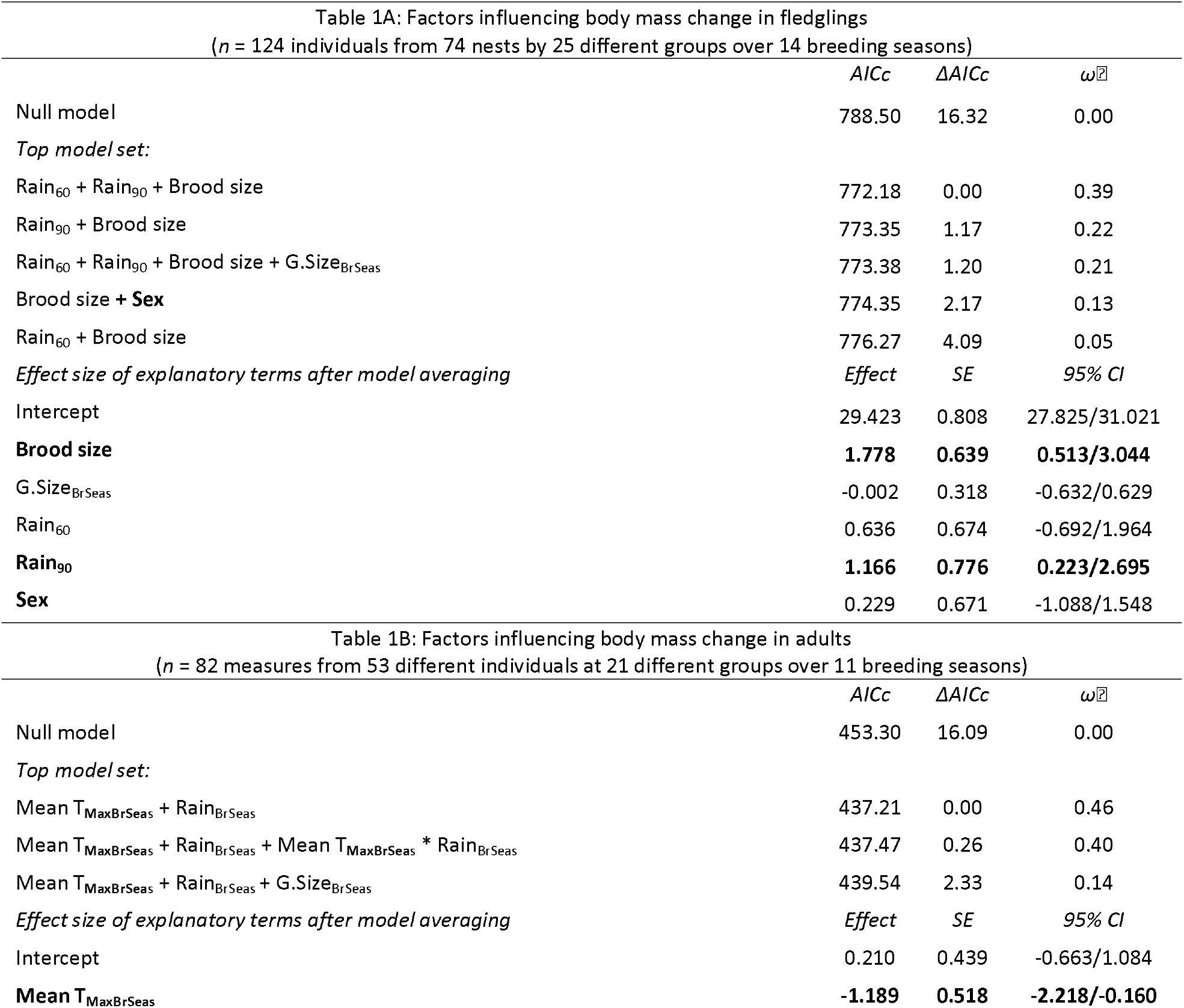
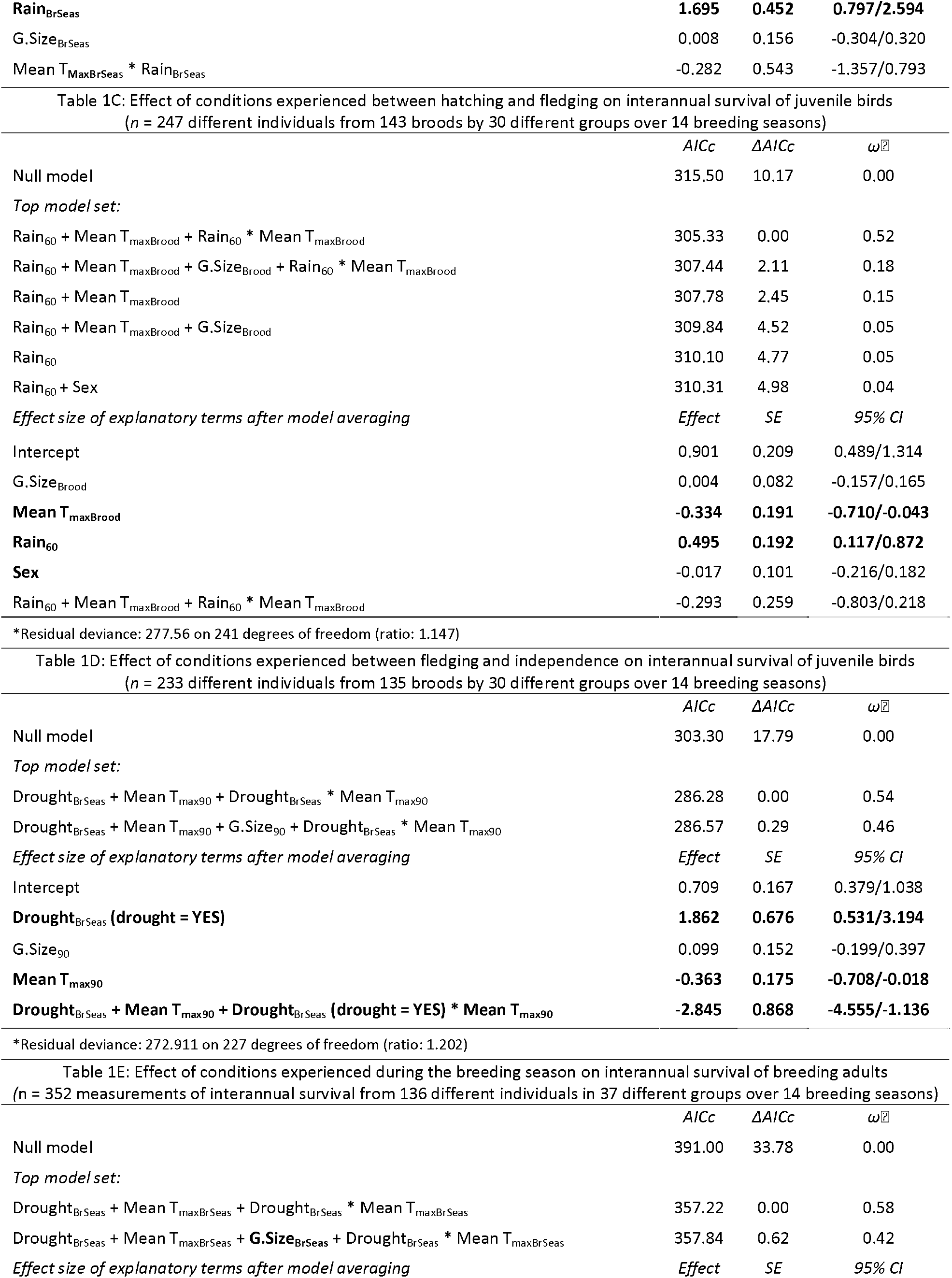
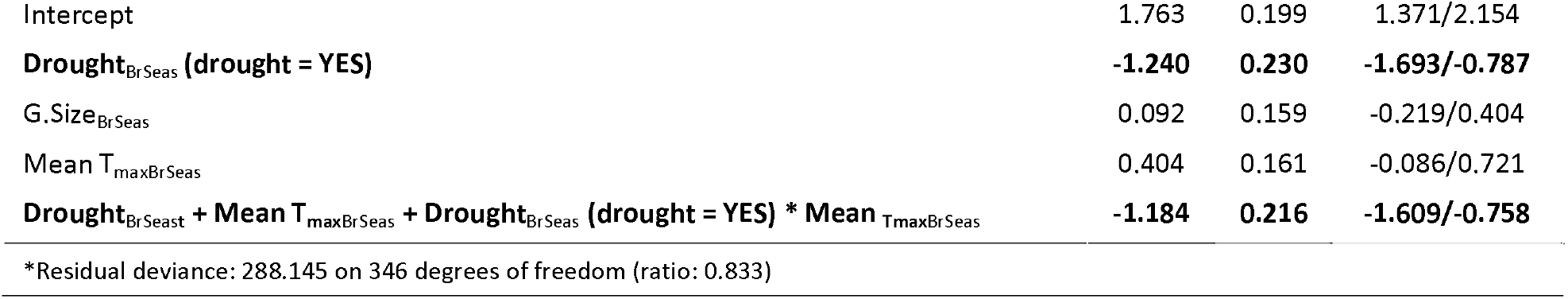
Top GLMM model sets for all analyses. Model averaging was implemented on all models with ΔAICc < 5. Significant terms after model averaging are shown in bold. Null models shown for comparison with top model sets. For full model sets, see Supporting Information.

**Fig 2:**
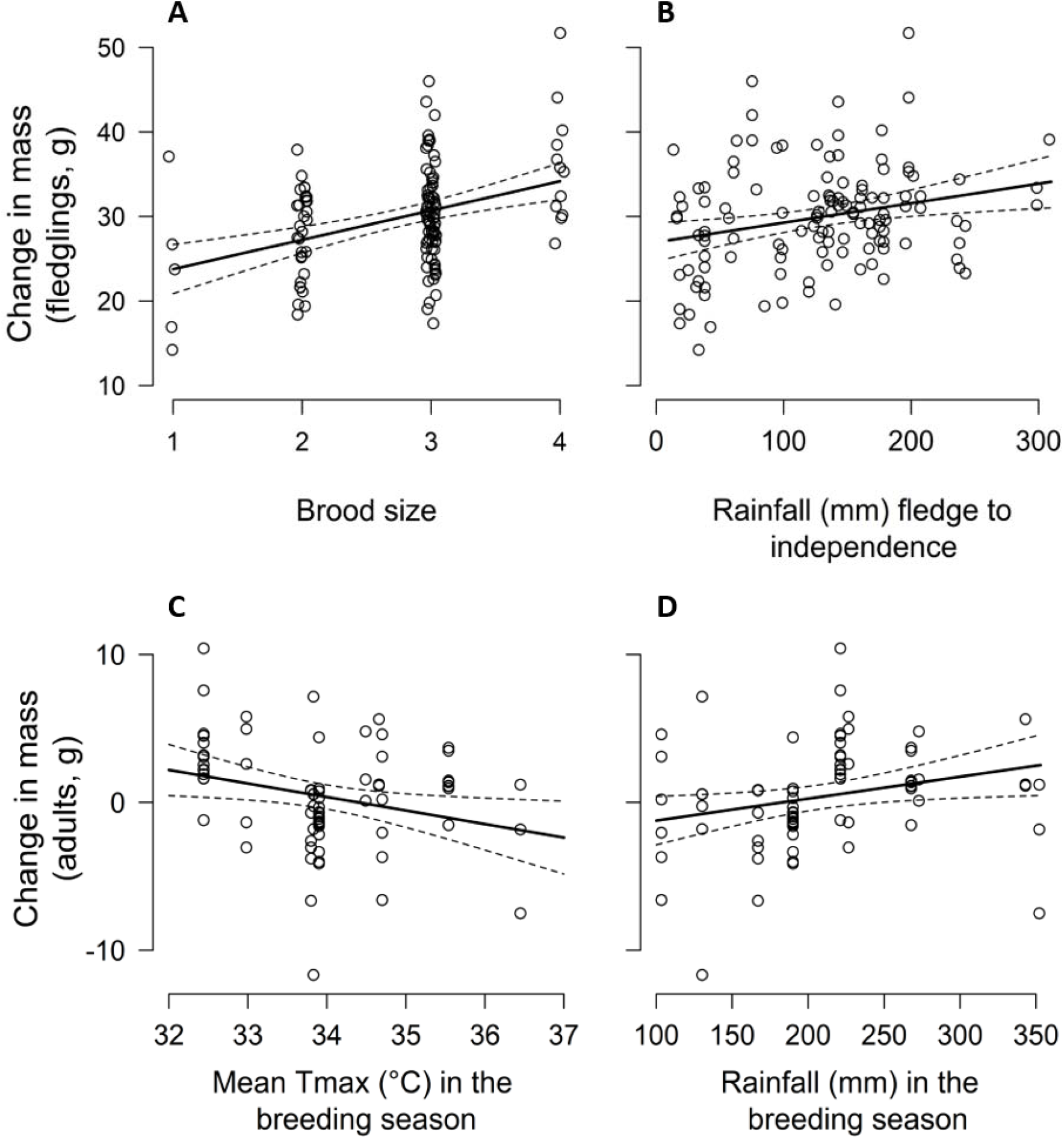
Change in body mass in southern pied babbler Turdoides bicolor fledglings between the fledging and nutritional independence in relation to (A) brood size and (B) rainfall (mm) between the fledging and nutritional independence and in adults between the start and end of the breeding season in relation to (C) average daily maximum temperatures and (D) total rainfall (mm) during the breeding season. Data points in (A) are jittered for improved visibility. Juvenile pied babblers from larger broods gained more mass between 11- and 90-days-of-age. This is likely explained by nestlings from larger broods fledgling at a smaller body mass, and hence growing more once they had left the nest, than those which fledged from smaller broods and may have attained larger mass while still in the nest.

High temperatures and low rainfall during the breeding season were associated with body mass loss between the start and the end of the breeding season in breeding adults (Table 1B; Fig. 2C, Fig. 2D). ΔM_b.Adults_ did not vary significantly with sex or group size, after model averaging, and nor did we find evidence of significant interactions among climate and group size variables (Table S2). Age was not associated with variation in ΔM_b.Adults_ in a subset of 36 individuals of known age (Table S3) and we therefore excluded age from the models analysing our full dataset presented here.

### Survival: juveniles

Of 596 nestlings of known Mass_11_, 254 (42.6%) survived to nutritional independence. Of these, 173 (68.1%) were present in the study population one year post-hatching. Natal group size ranged from 2–9 adults (mean = 4.4 ± 1.5). The likelihood of a juvenile surviving to one year of age increased as Rain_60_ increased (Table 1C, Fig. 3A). However, juveniles that experienced high mean T_maxBrood_ were less likely to survive to one year of age (Table 1C, Fig. 3B; see Table S4 for full model selection output).

**Fig 3:**
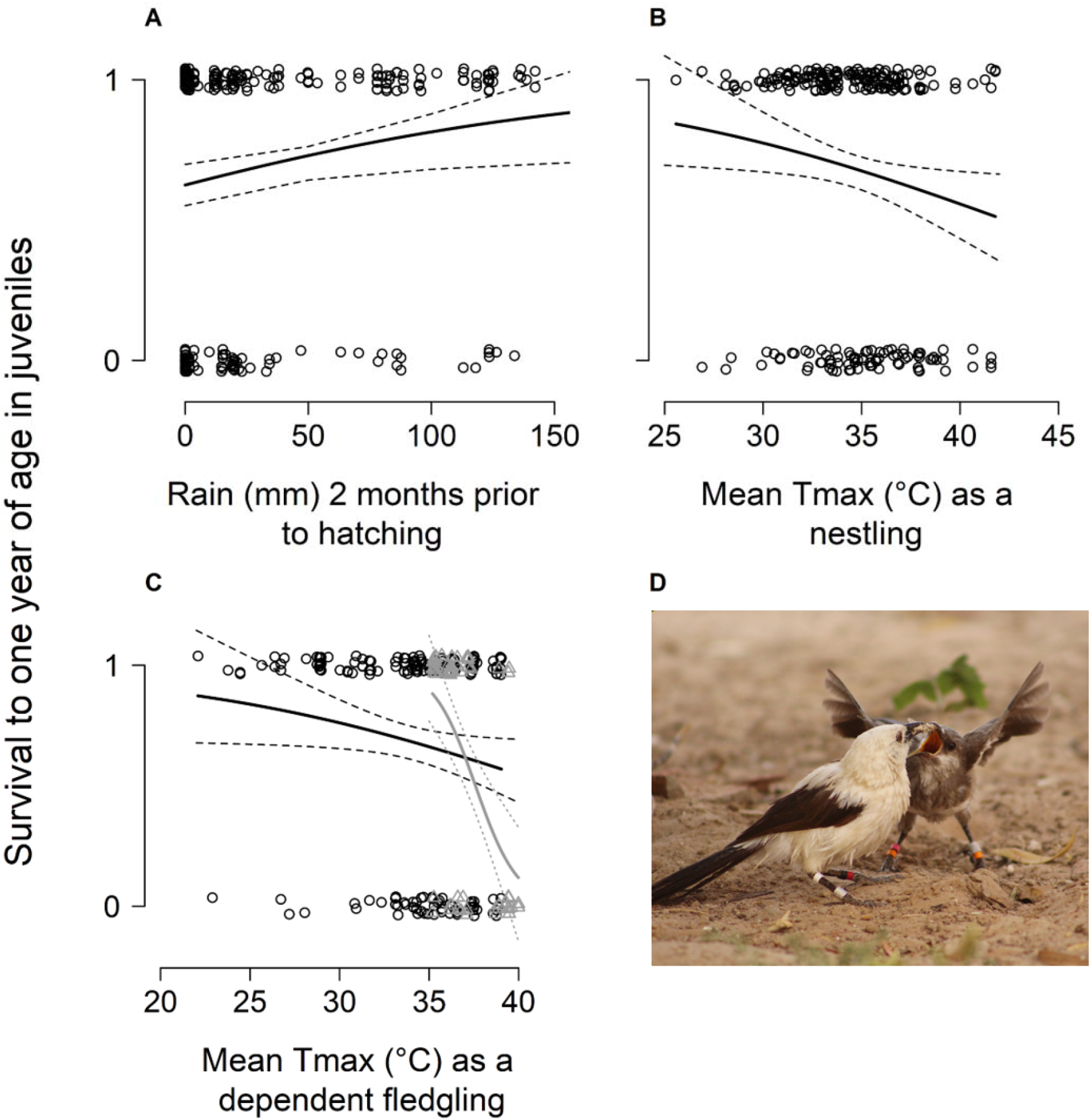
Interannual survival (0 = not present, 1 = present) in juvenile southern pied babblers Turdoides bicolor in relation to (A) rainfall prior to hatching; (B) mean daily maximum temperature between hatching and fledging; and (C) the interaction between temperature and drought between fledge and independence (non-drought breeding seasons: open circles, dashed confidence intervals, black colour; drought breeding seasons: open triangles, dotted confidence intervals, gray colour). Data points are integers (0,1) jittered for improved visibility. Individuals in the study population are uniquely identifiable by their colour and metal ring combinations (D; photograph by Nicholas B. Pattinson).

Juveniles were also less likely to survive to one year of age when they were exposed to high mean T_max90_ and Drought_BrSeas_. The effect of Drought_BrSeas_ on juvenile survival to one year of age was influenced by temperature: mean probability of survival was high (0.90 ± 0.05) when juvenile birds experienced both Drought_BrSeas_ and relatively cool mean T_max90_, whereas mean probability of survival was very low (0.12 ± 0.08) when juvenile birds experienced both Drought_BrSeas_ and high mean T_max90_ (Table 1D, Fig. 3C). This represents a more than seven-fold decrease in survival when individuals experienced both drought and high temperatures as dependent fledglings, compared to when drought occurred but temperatures were mild. Survival to one year of age did not vary significantly with Mass_11_, sex, brood size, or group size in either analysis. Survival to one year of age was also not significantly influenced by interactions between group size and environmental conditions (see Tables S4–S5 for full model selection output). Survival to one year of age was not associated with ΔM_b.Juv_ (GLMM: Est = 0.041 ± 0.037, *z* = 1.106, 95% CI: −0.039, 0.081).

### Survival: breeding adults

In 264 out of 352 records of interannual survival (75%; from 136 different individuals), breeding adults were still present at the start of the next breeding season. Breeding adults were less likely to be present at the start of the next breeding season when they had experienced Drought_BrSeas_ (Table 1E). The effect of Drought_BrSeas_ on the survival of breeding adults was influenced by temperature: mean probability of survival was high (0.81 ± 0.06) when individuals had experienced Drought_BrSeas_ alongside relatively cool mean T_maxBrSeas_. However, mean probability of survival was low (0.32 ± 0.09; Table 1E, Fig. 4) when individuals experienced both drought conditions alongside high mean T_maxBrSeas_, representing a more than 50% decrease in survival of breeding adults from one year to the next compared to when drought occurred but temperatures were mild. Interannual survival of breeding adults did not vary significantly with sex or group size, nor did we find evidence of significant interactions among climate and group size variables (see Table S6 for full model selection output). Age was not associated with variation in survival in a subset of 58 individuals of known age (Table S7) and we therefore excluded age from the models analysing our full dataset presented here. The probability of breeding adults surviving to the start of the next breeding season was not associated with ΔM_b.Adults_ (GLMM, Est = 0.031 ± 0.071, *z* = 0.436, 95% CI: −0.111, 0.171).

**Fig 4:**
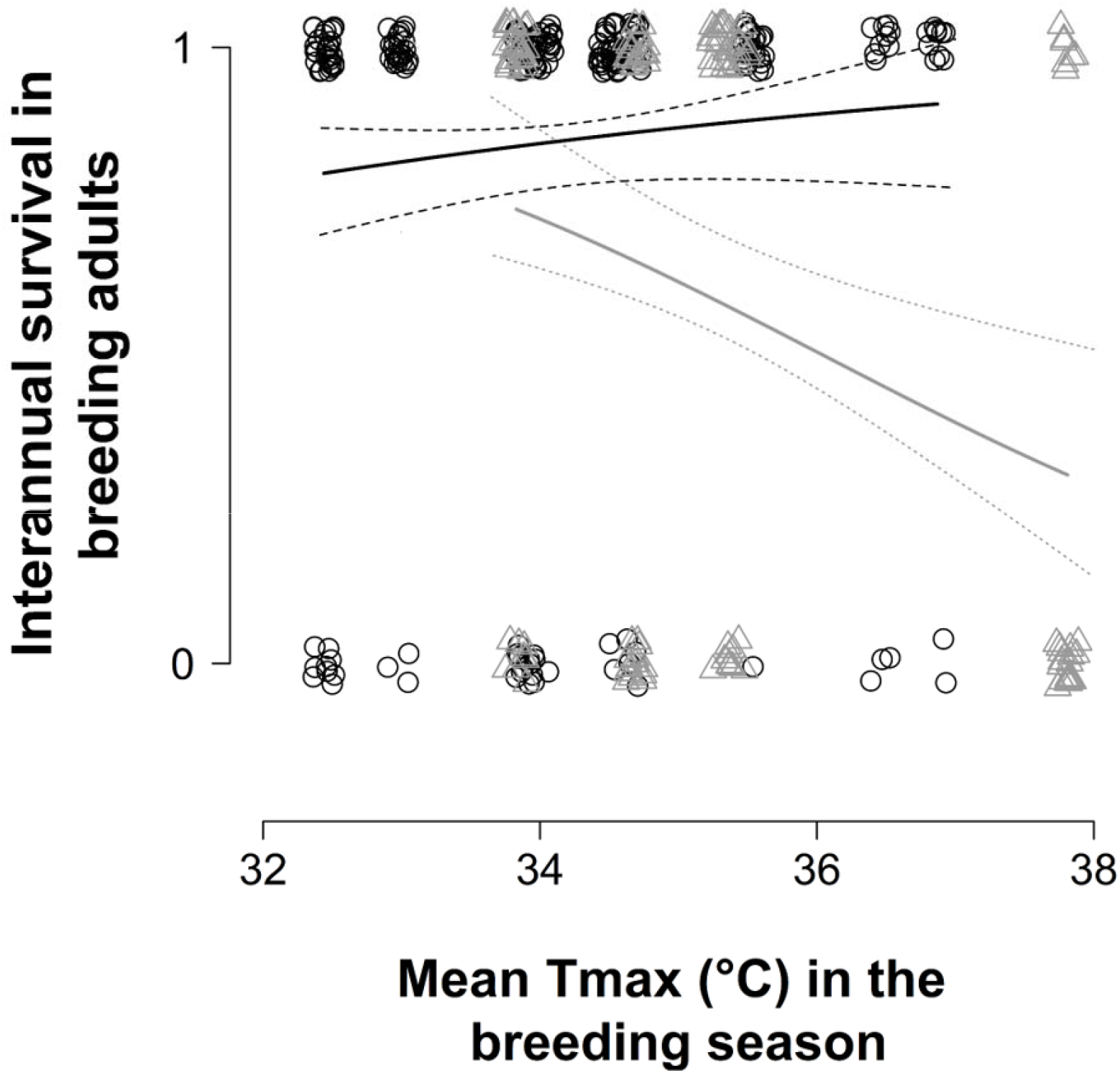
Survival of breeding adult southern pied babblers Turdoides bicolor from one breeding season to the next in relation to temperature and drought conditions during the initial breeding season (non-drought years: open circles, dashed confidence intervals, black colour; drought years: open triangles, dotted confidence intervals, gray colour). Data points are integers (0,1) jittered for improved visibility.

## Discussion

We investigated the potential for group size to buffer against the impacts of climatic factors on ΔM_b_ and interannual survival in a cooperatively breeding bird. We found that measures of ΔM_b_ and survival for individuals in larger groups were not affected differently by high temperatures and drought compared to those in smaller groups. Environmental conditions significantly affected ΔM_b_ and interannual survival in both juveniles and breeding adults across all group sizes, a finding that contributes to a rapidly growing body of literature (Overpeck 2013; Allen *et al.* 2015; Cruz-McDonnell & Wolf 2016) demonstrating that high temperatures and prolonged periods of low rainfall negatively impact on survival in a range of species.

Exposure to high temperatures and low rainfall was strongly associated with reduced growth in juvenile pied babblers and body mass loss in breeding adults. Body mass performs well as an index of condition (Labocha & Hayes 2012), particularly when change *within* individuals is measured over time, and poor body condition has been linked to reduced survival in both adult birds (e.g. Gardner *et al.* 2018) and nestlings (e.g. Todd *et al.* 2003; Schwagmeyer & Mock 2008). In our study population, in a simple univariate assessment, the within-individual ΔM_b_ we recorded did not appear to be significantly associated with interannual survival in either juvenile or breeding adult pied babblers, at least over our measurement windows, suggesting that reduced survival is not a consequence of body mass loss/reduced growth.

In pied babblers, exposure to chronic, sublethal effects of high temperatures and low rainfall within the same breeding season are associated with increased risk of overwinter mortality. Hot and dry conditions experienced by juvenile pied babblers between fledging and independence were associated with lower probability of survival to one year of age. Lower survival probabilities result in reduced recruitment into the adult population. Hot and dry conditions were also associated with strongly reduced likelihood of interannual survival in breeding adults. Together these processes likely contribute to the overall trend for population decline in below-average rainfall years in this species (Wiley 2017). Negative effects of adverse climate conditions on the interannual survival of breeding adults are particularly concerning because interannual survival rates of breeding adults (compared to non-breeding adult helpers) have the greatest impact on population growth rates in this species (Wiley 2017), and hence the probability of population persistence through time (Layton-Matthews *et al.* 2018).

Droughts are currently a regular, natural feature of the local climate (Tokura *et al.* 2018). Temperatures in the region have increased in recent decades (van Wilgen *et al.* 2016) and will continue to do so (IPCC 2013), and therefore an increase in the frequency of hot droughts can be expected. If this occurs, pied babbler populations are likely to decline as altered drought regimes reduce opportunities for population recovery following hot drought events. Consecutive years characterised by exceptionally hot and dry conditions could lead to failed recruitment and population crashes, as has been observed in *Athene cunicularia* burrowing owls (Cruz-McDonnell & Wolf 2016) and is predicted for pied babblers (Wiley 2017; Conradie *et al.* 2019).

Group size did not buffer the impacts of hot, dry weather on individual body mass change or survival in juveniles or breeding adults, since birds across all group sizes were similarly affected. This is consistent with the findings of two concurrent studies on a cooperatively-breeding mammal (van de Ven *et al.* 2019a) and a cooperatively breeding bird (Guindre-Parker & Rubenstein 2020). Adverse effects of climatic conditions on ΔM_b_ and survival are therefore likely driven primarily by physiological tolerance limits (Smit *et al.* 2018) and resource constraints (Nowakowski *et al.* 2018) acting on individuals, irrespective of the number of individuals present in their social group. Benefits of group living and cooperation, including load-lightening and the production of more surviving young by larger groups, have been observed in pied babblers previously (Ridley 2016; Wiley & Ridley 2016; Bourne *et al.* 2020). Yet we show here that larger group sizes did not moderate the influence of high temperatures and drought on body mass change or survival. While it is possible that the benefits of larger group sizes and the presence of helpers may have previously helped to mitigate the effects of adverse environmental conditions (Jetz & Rubenstein 2011; Russell 2016; Cornwallis *et al.* 2017), it appears that any such advantage is no longer detectible given current extreme conditions and a rapidly changing climate. Buffering effects of group size may be detectable in reproductive outputs rather than measures of mass change and survival, although see Bourne et al. (n.d.) which suggests that this is not the case in pied babblers.

The Intergovernmental Panel on Climate Change predicts that the incidence of hot and dry extremes will continue to become more frequent over most land masses (IPCC 2013). In arid and semi-arid regions already affected by both decreased precipitation and increased warming, interannual survival and recruitment in resident avian species (such as this population of pied babblers) may be insufficient to allow for population recovery between hot droughts. Taken together with our finding that larger group sizes did not buffer pied babblers against adverse climatic conditions, our data raise concerns for the long-term persistence of arid-zone species in the face of changing climatic conditions. The adaptive benefits of cooperative life history strategies in highly variable environments are unlikely to be sufficient to counteract the impacts of rapidly changing climatic conditions, in particular the increased frequency of climate extremes such as heat waves and drought.

## Acknowledgements

We thank the management teams at the Kuruman River Reserve (KRR) and surrounding farms, Van Zylsrus, South Africa, for making the work possible. The KRR was financed by the Universities of Cambridge and Zurich, the MAVA Foundation, and the European Research Council (Grant No. 294494 to Tim Clutton-Brock), and received logistical support from the Mammal Research Institute of the University of Pretoria. We thank Sello Matjee, Paige Ezzey, and Lesedi Moagi for fieldwork assistance during 2016–2019, and all past and present staff and students of the Pied Babbler Research Project for data collected since 2003. Dr Todd Erickson provided valuable comments on an early draft. This work was funded by the DST-NRF Centre of Excellence at the FitzPatrick Institute for African Ornithology, the University of Cape Town, the Oppenheimer Memorial Trust (Grant No. 20747/01 to ARB), the British Ornithologists’ Union, the Australian Research Council (Grant No. FT110100188 to ARR), a BBSRC David Phillips Fellowship (BB/J014109/1 to CNS), and the National Research Foundation of South Africa (Grant No. 110506 to SJC). The opinions, findings and conclusions are those of the authors alone, and the National Research Foundation accepts no liability whatsoever in this regard. We thank the three knowledgeable anonymous reviewers for their constructive comments which helped to improve this work tremendously.

